# Mining the Selective Remodeling of DNA Methylation in Promoter Regions to Identify Robust Gene-Level Associations with Phenotype

**DOI:** 10.1101/2020.01.05.895326

**Authors:** Yuan Quan, Fengji Liang, Yuexing Zhu, Ying Chen, Ruifeng Xu, Jianghui Xiong

**Affiliations:** School of Computer Science and Technology, Harbin Institute of Technology Shenzhen Graduate School, Shenzhen, Guangdong, 518055, China; Lab of Epigenetics and Advanced Health Technology, Space Institute of Southern China, Shenzhen, Guangdong, 518117, China; State Key Laboratory of Space Medicine Fundamentals and Application, China Astronaut Research and Training Center, Beijing, 100094, China; Aromability Inc., Beijing, 100080, China

**Keywords:** DNA methylation, remodeling, gene level, SIMPO algorithm, phenotype-associated genes

## Abstract

Epigenetics is an essential biological frontier linking genetics to the environment, where DNA methylation is one of the most studied epigenetic events. In recent years, through the epigenome-wide association study (EWAS), researchers have identified thousands of phenotype-related methylation sites. However, the overlap between identified phenotype-related DNA methylation sites are often quite small, and it might clue to methylation remodeling has a certain degree of randomness within the genome. Thus, the identification of robust gene-phenotype associations is crucial for interpreting pathogenesis. How to integrate the methylation values of different sites on the same gene and to mining the DNA methylation at the gene level remains a challenge. A recent study found that the DNA methylation difference of the gene body and promoter region has a strong correlation with gene expression. In this study, we proposed a Statistical difference of DNA Methylation between Promoter and Other Body Region (SIMPO) algorithm to extract DNA methylation values at the gene level. First, by choosing to smoke as an environmental exposure factor, our method led to significant improvements in gene overlaps (from 5% to 17%) between different datasets. In addition, the biological significance of these genes (∼23%) are significantly better than those identified by traditional probe-based methods (∼18%, P-value = 5.18e-03). Then, we selected two disease content (e.g., insulin resistance and Parkinson’s disease) to show that the biological efficiency of disease-related gene identification increased from 15.43% to 44.44% (P-value = 1.20e-28). Thus, our results declare that mining the selective remodeling of DNA methylation in promoter regions can identify robust gene-level associations with phenotype, and the characteristic remodeling of a given gene’s promoter region can reflect the essence of disease.

## 1. Introduction

Epigenetics is a branch of genetics that studies the heritable changes in gene expression without changing the nucleotide sequence of a gene [1], including DNA methylation, histone modification, and regulation of non-coding RNA, among which DNA methylation is currently a hot topic in epigenetics [2]. Several studies have shown that the regulation of genes by DNA methylation is related to the occurrence and development of various diseases, such as cancer [3-5], cardiovascular and cerebrovascular diseases [6-8], and metabolic diseases [9,10].

Similar to GWAS (Genome-wide association study), EWAS compares the variation between patients and healthy people at the DNA methylation level and associates epigenetic variation with complex diseases as well as interprets the pathogenic cause of complex diseases at the epigenetic level [11]. EWAS opens the door for researchers to study complex diseases, allowing us to find several previously undiscovered disease-related methylation sites, providing more epigenetic mechanisms for the pathogenesis of complex diseases [12,13]. Since 2009 when the first EWAS study was published, EWAS research has grown exponentially in recent years, reaching 618 publications in 2019 [12]. Due to the availability of whole blood DNA methylation data, most of the materials for most current EWAS studies are focused on whole blood tissues [12].

In the detection of clinical samples, the human DNA methylation chip is a common method for high-throughput EWAS analysis. The currently widely used methylation chip is the Illumina 450K BeadChip [11-13]. However, multiple methylation probes are distributed in the same functional region of the same gene in the 450K BeadChip, and different probes will detect different methylation values. In addition, because the methylation modification has a certain degree of randomness on the genome, the results of the similar EWAS studies are often inconsistent [14]. For example, there are several EWASs that focus on smoking-related phenotypes and identify tens of thousands of significantly different probes [15-22]. We found that starting from these different probes, each independent EWAS can correspond to 101∼6,180 differential genes, and these EWAS publications predicted a total of 7,340 genes. However, only 1,334 (18.17%) of genes were present in two or more independent EWAS. In addition, four diabetes-related EWAS projects predicted 493 [23], 565 [24], 1,179 [25], and 3,186 [26] diabetes-related genes, respectively. However, only 7.82% (392 of 5,012 genes) of these genes were simultaneously identified in multiple EWASs.

Another example is the identification of Parkinson’s disease-related 162 significantly different DNA methylation probes based on three EWASs, corresponding to 194 genes [27-29]. Unfortunately, the intersection of only one gene, *STK38L*, existed in these three studies. Therefore, traditional probe-based EWASs have some limitations in identifying phenotype-related genes based on differential probes. Moreover, how to integrate the DNA methylation values of different probes on the same gene and characterize the DNA methylation degree at the gene level has become a challenge for traditional EWASs.

Because methylation remodeling has a certain degree of randomness and complexity on the genome, there is no significant correlation to only consider the remodeling of DNA methylation in promoter regions or to only consider the remodeling of DNA methylation in body regions with gene expression (**Figure 1**). Therefore, this study proposed that by combining the DNA methylation remodeling of the promoter regions and the body regions, we could identify robust gene-level associations with the phenotype (**Figure 1**). Recently, researchers noticed that the correlation between DNA methylation difference of the gene body and promoter region (MeGDP) and gene expression is up to 0.67 [30]. This result is consistent with our conjecture and suggests that DNA methylation differences between the gene promoter and body regions can be used as a DNA methylation index to predict gene expression (**Figure 1**).

**Figure 1.**
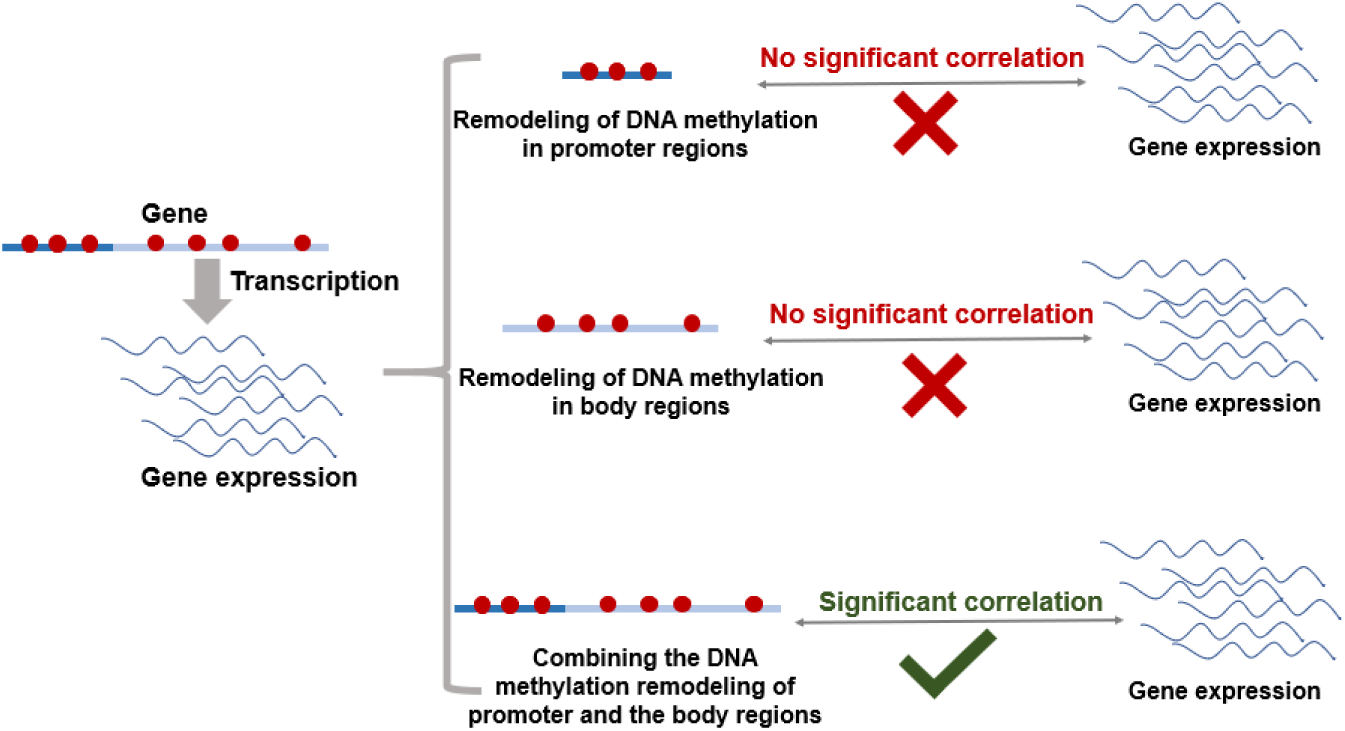
The correlations of DNA methylation remodeling in promoter and body regions with gene expression. Dark blue lines represent the promoter regions of the gene, and light blue lines represent the body regions of the gene. Red dots represent DNA methylation sites on the genome.

Based on the above correlation, this study proposed the statistical difference of DNA Methylation between Promoter and Other Body Region (SIMPO) algorithm to mining gene-level DNA methylation associations with phenotype. It showed the robustness of SIMPO-identified differential genes in the same dataset and between different datasets through three smoking phenotype-related DNA methylation datasets. The results also showed that the biological significance of SIMPO-identified differential genes are significantly better than those predicted by traditional probe-based methods. In addition, we further applied the SIMPO algorithm to predict insulin resistance (IR)- and Parkinson’s disease (PD)-associated genes and revealed the biological significance of corresponding genes.

## 2. Materials and Methods

### 2.1 The DNA methylation and transcription data collection

First, this study collected transcription and DNA methylation data from the MESA (Multi-Ethnic Study of Atherosclerosis) Epigenomics and Transcriptomics Study. This study has been launched to investigate potential gene expression regulatory methylation sites in humans by examining the association between CpG methylation and gene expression in purified human monocytes from 1,202 individuals (ranging 44∼83 years of age) and proved that blood monocyte transcriptome and epigenome can reveal loci associated with human age [31]. We downloaded the above data from the NCBI GEO (Gene Expression Omnibus) database (GEO accession: GSE56045 and GSE56046).

Next, we used three smoking phenotype-related DNA methylation datasets to test the robustness of the SIMPO algorithm. Previous studies have found that smoking can cause a variety of smoking-related diseases by affecting DNA methylation and causing abnormal gene expression [20,32,33]. For example, based on peripheral blood DNA methylation data of 464 individuals who were current, former, and never-smokers (GEO accession: GSE50660), researchers have identified 15 methylation sites associated with smoking [32]. In addition, the GSE53045 dataset contains DNA methylation data extracted from the peripheral mononuclear cell of 50 smokers and 61 non-smokers. Moreover, 910 significant loci have predicted after Benjamini-Hochberg correction based on this dataset [20]. The third smoking phenotype-related DNA methylation dataset was collected from GSE85210. GSE85210, including DNA methylation data of 172 smokers and 81 non-smokers blood cells, and revealed 738 CpGs significantly associated with current smoking [33].

Data of IR-related DNA methylation BeadChip analyzed in this study were downloaded from the NCBI GEO database (GEO accession: GSE115278). This dataset uses Illumina HumanMethylation450 BeadChip’s GPL16304 platform and contains DNA methylation data of peripheral white blood cells collected from 74 HOMA-IR>3, and 258 HOMA-IR≤3 individuals. Furthermore, based on this data, a rigorous statistical analysis revealed that 478 CpGs showed a differential methylation pattern between individuals with HOMA-IR□≤□3 and >□3 [34].

Two PD-related DNA methylation datasets were downloaded from the NCBI GEO database (GEO accession: GSE72774 and GSE111629). These dataset uses Illumina HumanMethylation450 BeadChip’s GPL13534 platform. GSE72774 contains DNA methylation data of whole blood collected from 289 individuals with PD and 219 control samples then, these researchers obtained 82 genome-wide significant CpGs of PD [27]. Whole blood DNA methylation data of GSE111629 were collected from 335 PD individuals and 237 controls.

### 2.2 SIMPO algorithm

Because previous research found that the DNA methylation difference between the promoter region and the body region is highly related to the expression level of the gene [15]. The input data of the SIMPO algorithm is the DNA methylation beta value of cg probes that are located in the promoter regions and the other regions. The statistical difference method *t*-test is used to SIMPO, and the degree of difference (SIMPO score) is used to characterize the DNA methylation of corresponding genes:

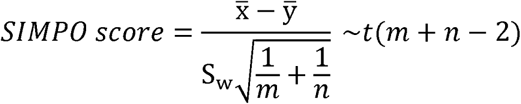

where

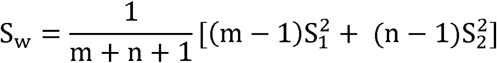

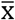: average DNA methylation value of all probes that are located in the other region; 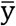: average DNA methylation value of all probes that are located in the promoter region; m: number of probes that are located in the other region; n: number of probes that are located in the promoter region; 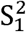: variance of DNA methylation values of probes that are located in the other region; 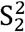: variance of DNA methylation values of probes that are located in the promoter region. In addition, since the SIMPO score relates to the number of probes, in order to ensure the reliability of the SIMPO score, we only selected genes with several other region-located and promoter region-located probes greater than or equal to five for further calculation.

### 2.3 Disease-associated genes collection

In this study, disease-associated genes were collected from the DisGeNET database (http://www.disgenet.org) and the SCG-Drug database (http://zhanglab.hzau.edu.cn/scgdrug) [35,36]. DisGeNET database integrates multiple disease gene databases, and gene-disease associations (GADs) reported in a large number of works of literature. Data sources include UniProt, Comparative Toxicogenomics Database (CTD), ClinVar, Orphanet, GWAS Catalog, Genetic Association Database. The latest version is v5.0, which contains 561,119 gene-disease pairs involving 17,074 genes and 20,370 diseases. In addition, DisGeNET v5.0 has developed a gene-disease relationship scoring model with scores between 0 and 1. Higher scores indicate higher confidence in the gene-disease relationship [35]. DisGeNET score for the gene-disease relationship is supported by multiple pieces of evidence and has high confidence. The SCG-Drug database collects gene-disease associations from multiple sources [36]. Similar to DisGeNET, SCG-Drug also annotates the scoring model of gene-disease associations.

### 2.4 Noise generation

In order to verify the robustness of SIMPO algorithm in the same dataset, this study added random noise of 0.1 to 1 degree to the DNA methylation beta value of each probe. Firstly, because the range of normalized DNA methylation beta is −1 to 1, we generated the random numbers in the range of −1 to 1 through the Python random module. Next, we multiplied the random number by 0.1 to 1 and obtained the random noise values of about 0.1 to 1. Third, we added the random noise values to the original DNA methylation beta values and received the new beta values. We further used these new DNA methylation beta values of probes to the SIMPO score calculation.

### 2.5 KEGG pathway enrichment

We enriched the KEGG pathway of folic acid supplement-derived differential genes through GSEA (Gene Set Enrichment Analysis) [37]. The rank of differential genes was according to SIMPO scores of PDs, and KEGG pathway gene set was downloaded from Molecular Signatures Database (MSigDb, c2.cp.kegg.v6.2.symbols.gmt). GSEA calculations are performed based on the R packages of ‘dplyr’ and ‘GSEABase’.

## 3. Results

### 3.1 Correlation between SIMPO score and transcription value of gene

In this study, the DNA methylation feature (SIMPO score) of each gene was extracted based on the SIMPO algorithm, and the Spearman correlation test was used to test the correlation between the SIMPO score and mRNA transcription average of each gene in 1202 samples (GSE56045 and GSE56046 datasets). The results are shown in **Figure 3**: based on SIMPO-TSS200 algorithm, the SIMPO scores of 43.44% of the genes are significantly related to the average mRNA transcription (**Figure 3g**) (**Table S1**); based on SIMPO-TSS1500 algorithm, the SIMPO scores of 41.22% of the genes are significantly related to the average mRNA transcription (**Figure 3h**) (**Table S2**); based on SIMPO-TSS200&TSS1500 algorithm, the SIMPO scores of 41.18% genes are significantly correlated with the average mRNA transcription (**Figure 3i**) (**Table S3**). The above results are similar to the significant correlation ratio of probes based on DNA methylation beta value (**Figure 3a∼3f**). It is indicated that the SIMPO score of the gene has a good correlation with the average mRNA transcription, and the SIMPO score can contain the original DNA methylation information of the gene.

**Figure 2.**
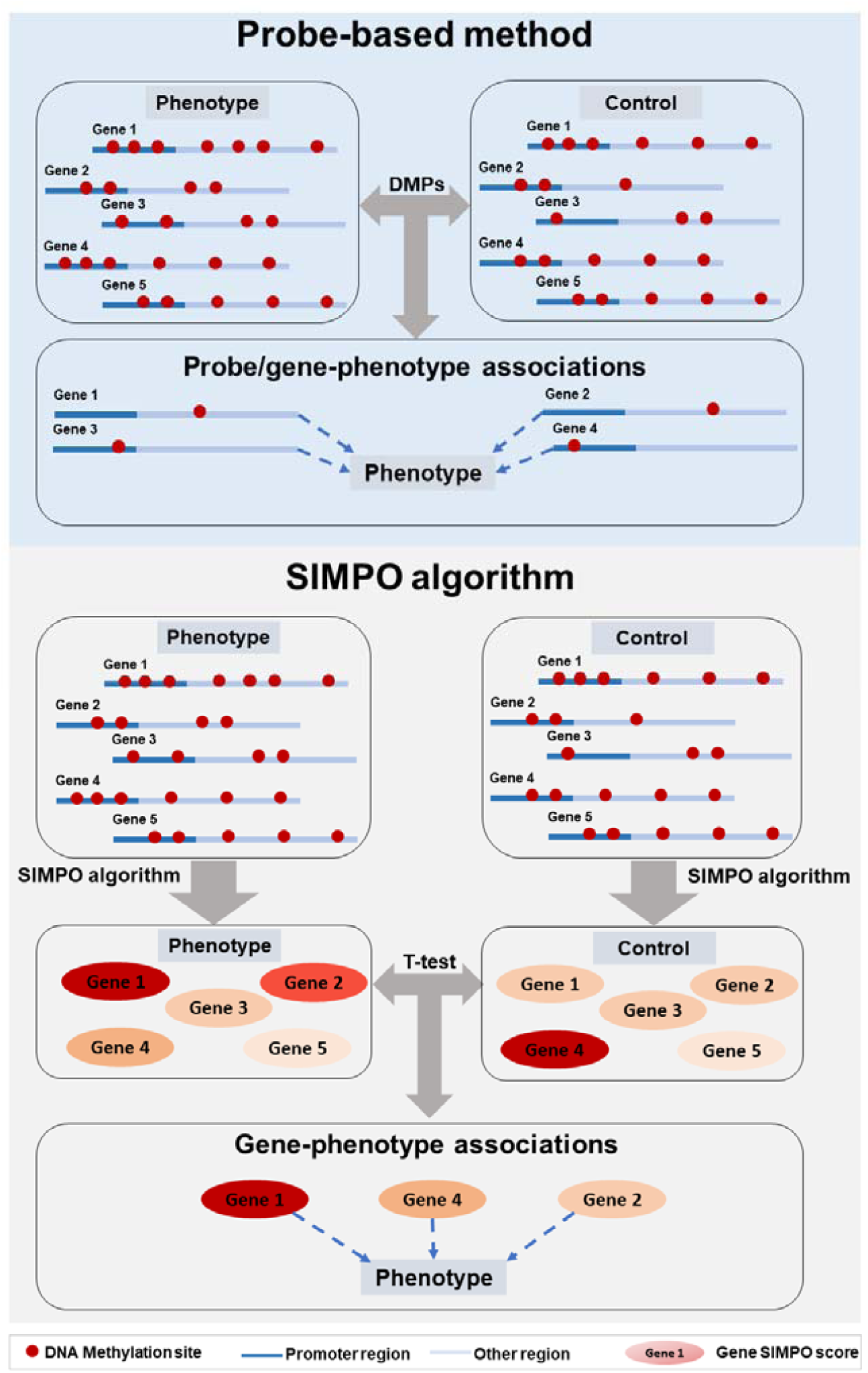
Pipeline comparison of probe-based method and SIMPO algorithm.

**Figure 3.**
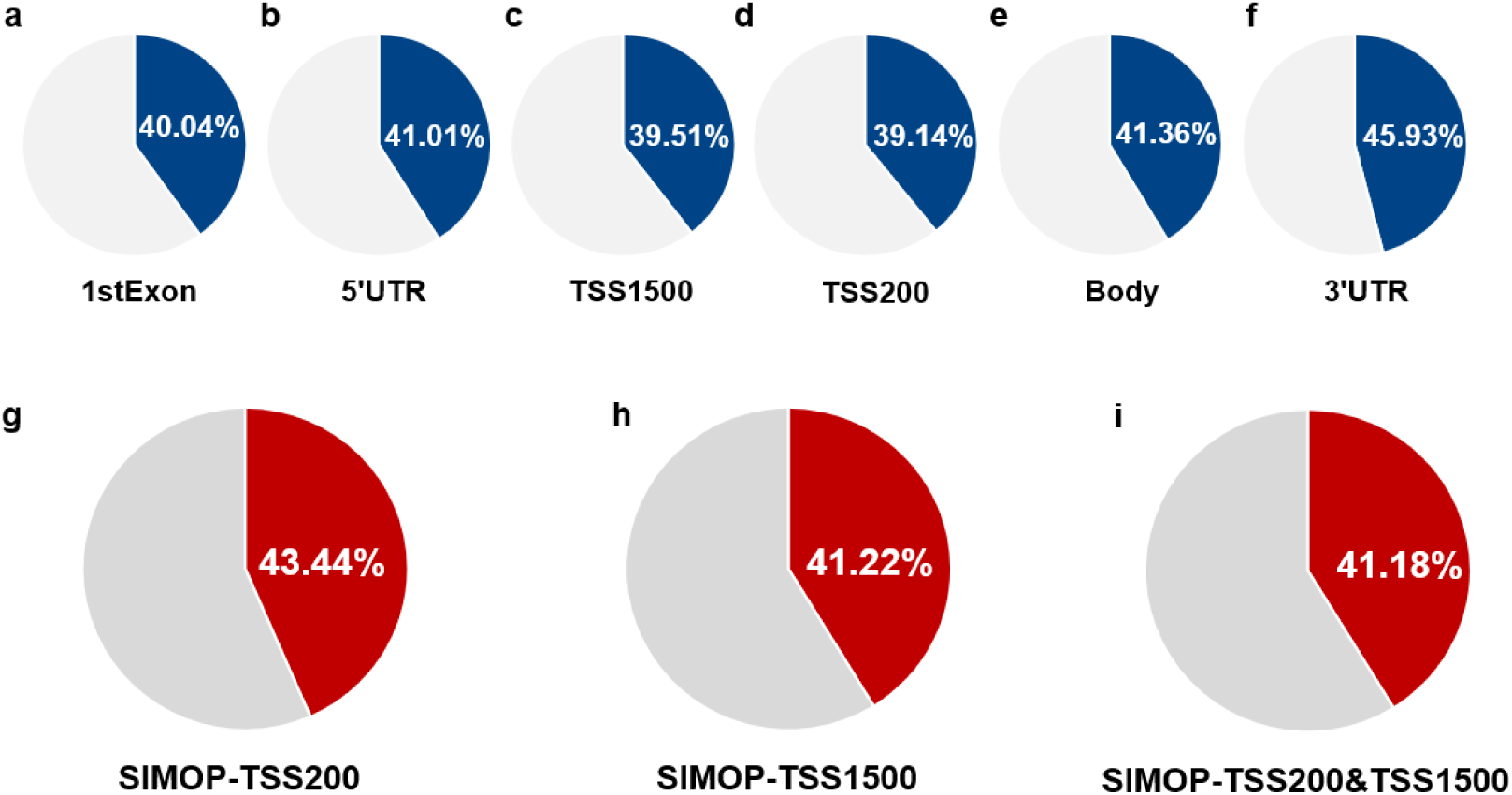
Proportion of probes/genes whose DNA methylation beta values or SIMPO scores are significantly associated with mRNA transcription values. Correlation analysis is based on the Spearman correlation test. (a) for probes located in 1stExon regions. (b) for probes located in 5’UTR regions. (c) for probes located in TSS1500 regions. (d) for probes located in TSS200 regions. (e) for probes located in body regions. (f) for probes located in 3’UTR regions. (g) for SIMPO-TSS200 algorithm-identified genes. (h) for SIMPO-TSS1500 algorithm-identified genes. (i) for SIMPO-TSS200&TSS1500 algorithm-identified genes.

### 3.2 Robustness verification of SIMPO algorithm for the same datasets

Based on the SIMPO algorithm and traditional probe-based algorithm, DNA methylation features of different genes of smokers and healthy people were obtained. Then, the significantly associated probes/genes of smoking were predicted through differential analysis (calculated by *t*-test). The number of differential genes (P-value ≤ 0.05) obtained from these three smoking DNA methylation datasets are shown in **Figure 4** (**Table S4-S8**).

**Figure 4.**
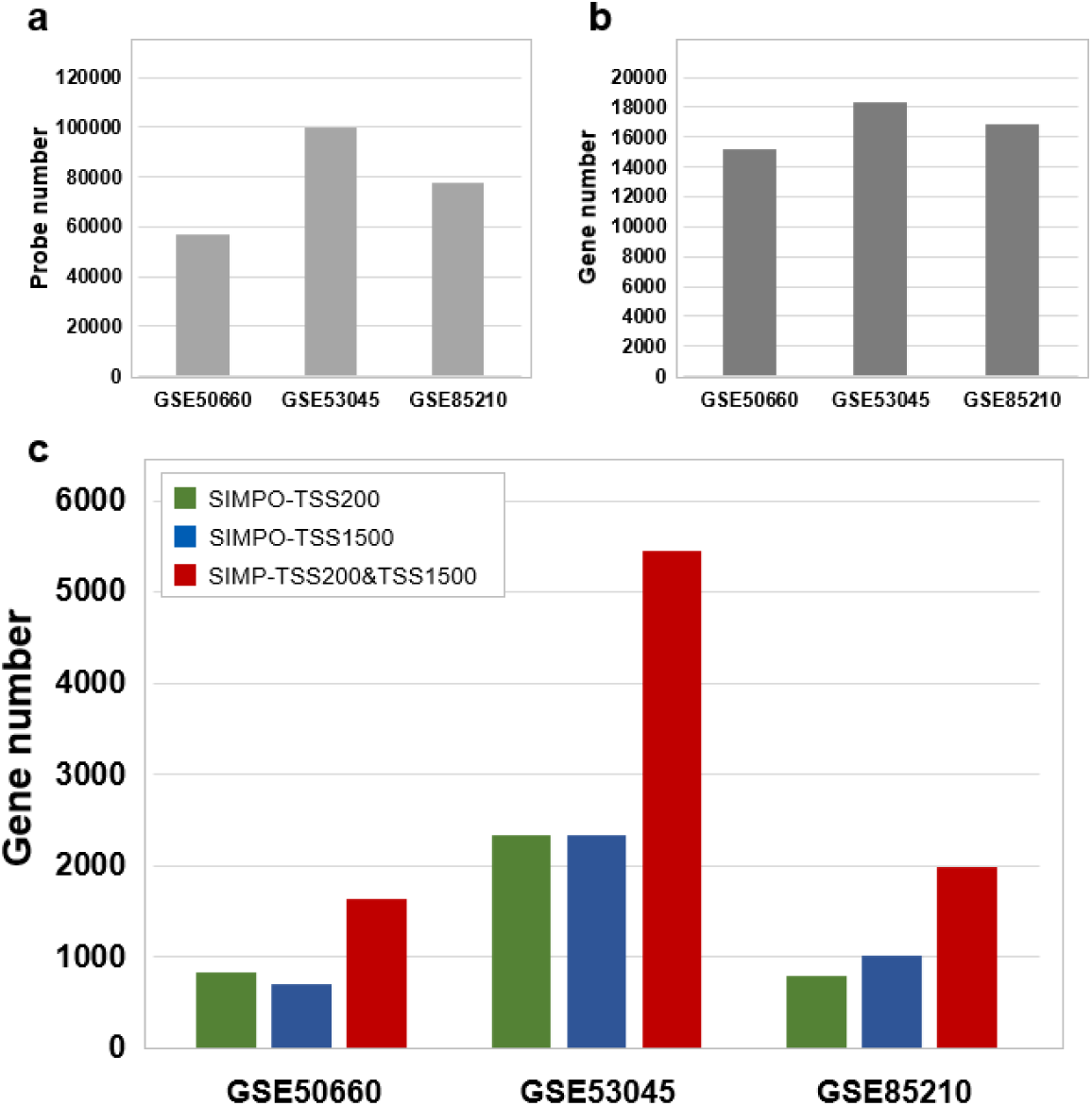
The significantly associated probe/gene numbers of smoking phenotype identified by different methods. (a) The significantly associated probe numbers of three smoking phenotype-related datasets identified by DMPs. (b) The significantly associated gene numbers of three smoking phenotype-related datasets identified by DMGs. (c) The significantly associated probe numbers of three smoking phenotype-related datasets identified by SIMPO algorithms.

For a particular gene, multiple probes contained in it will get different P-values. We selected the max P-value and the min P-value of the probe to represent the correlation between this gene and the smoking phenotype, and then obtained the ranking of these genes, respectively. We compared the intersection of the top N genes to show the robustness of traditional EWAS, which often focus only on the DNA methylation level of the probes for the same dataset. The results are shown in **Figure 5**: For the probe-focused study, the robustness in the same dataset is weak, and only about 8% of the genes have intersections.

**Figure 5.**
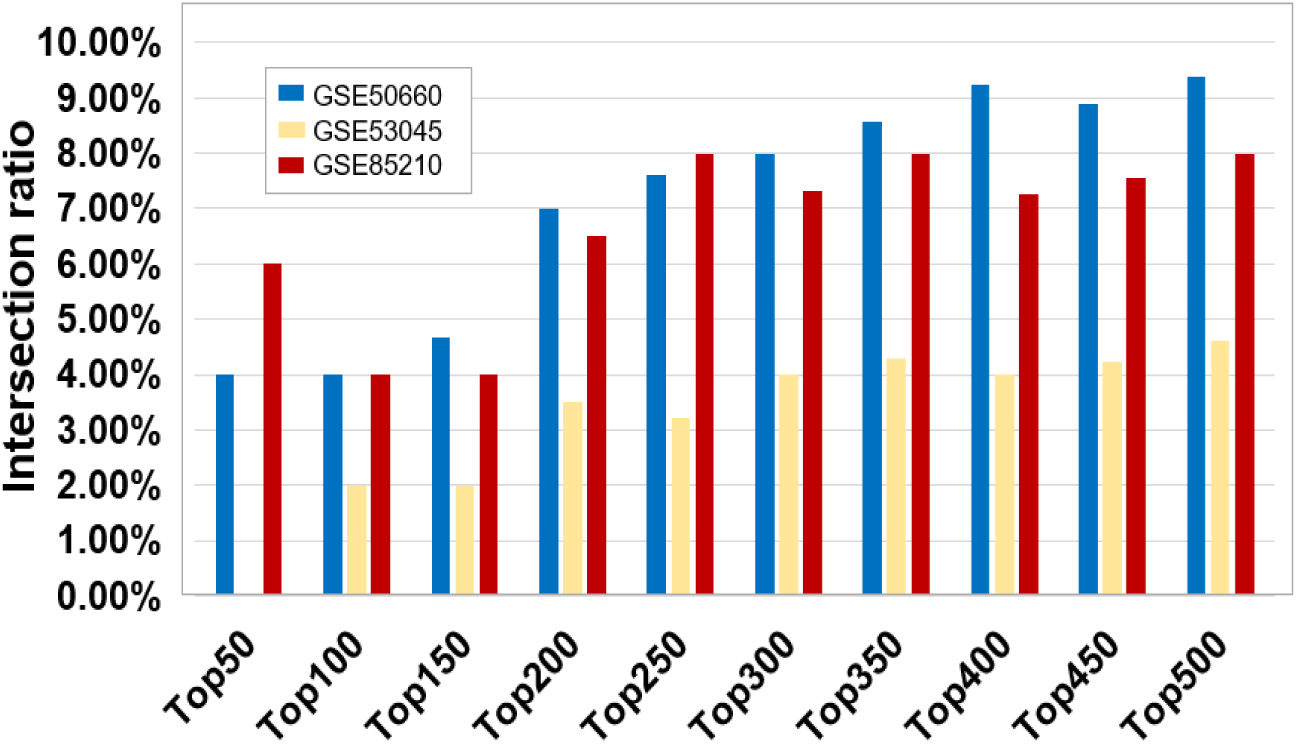
The intersection ratios of top N differential genes derived from a different probe located in the same genes (between the max P-value probe and min P-value probe).

Next, in order to test the robustness of the SIMPO algorithm in the same dataset, this study added random noise of 0.1 to 1 degree to the three DNA methylation data related to the smoking phenotype. Moreover, the intersections of top N smoking-associated genes identified by the original data and after adding noise-data between the traditional probe-based methods (DMPs and DMGs) and the SIMPO algorithm were compared. The results are shown in **Figures 6 and 7**: For the GSE50660 and GSE85210 datasets, when different levels of noise are added, the gene intersections obtained by the SIMPO algorithm were more significant than probe-based methods. Hence, the robustness of SIMPO is better than the traditional probe-based method for the same dataset.

**Figure 6.**
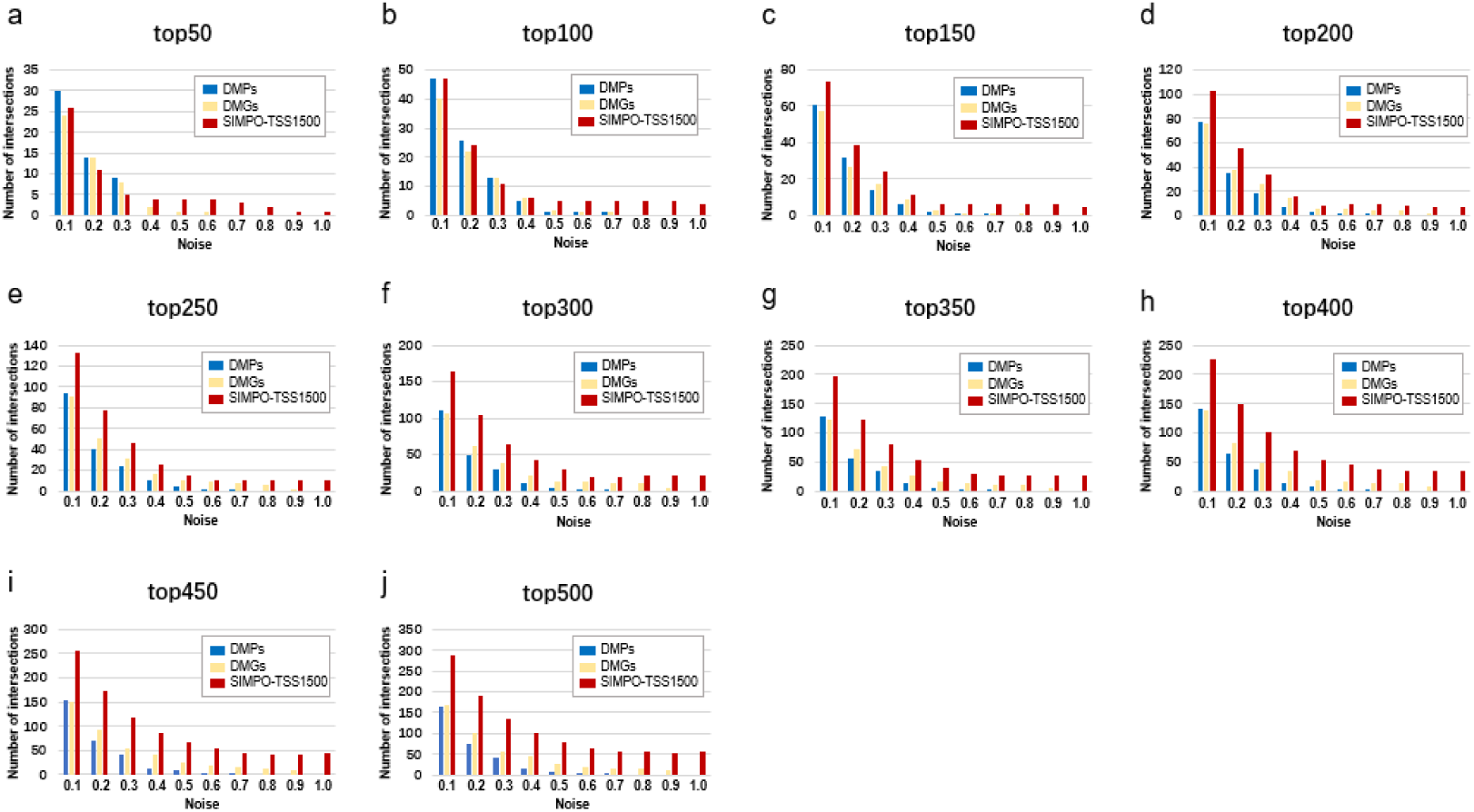
The intersections of top N smoking-associated genes identified by the original data and after adding noise-data of the GSE50660 dataset. The blue bars represent genes identified by the DMPs, the yellow bars represent genes identified by the DMGs, and the red bars represent genes identified by the SIMPO algorithm. (a) for to p50 probes/genes. (b) for top 100 probes/genes. (c) for top 150 probes/genes. (d) for top 200 probes/genes. (e) for top 250 probes/genes. (f) for top 300 probes/genes. (j) for top 350 probes/genes. (h) for top 400 probes/genes. (i) for top 450 probes/genes. (j) for top 500 probes/genes.

**Figure 7.**
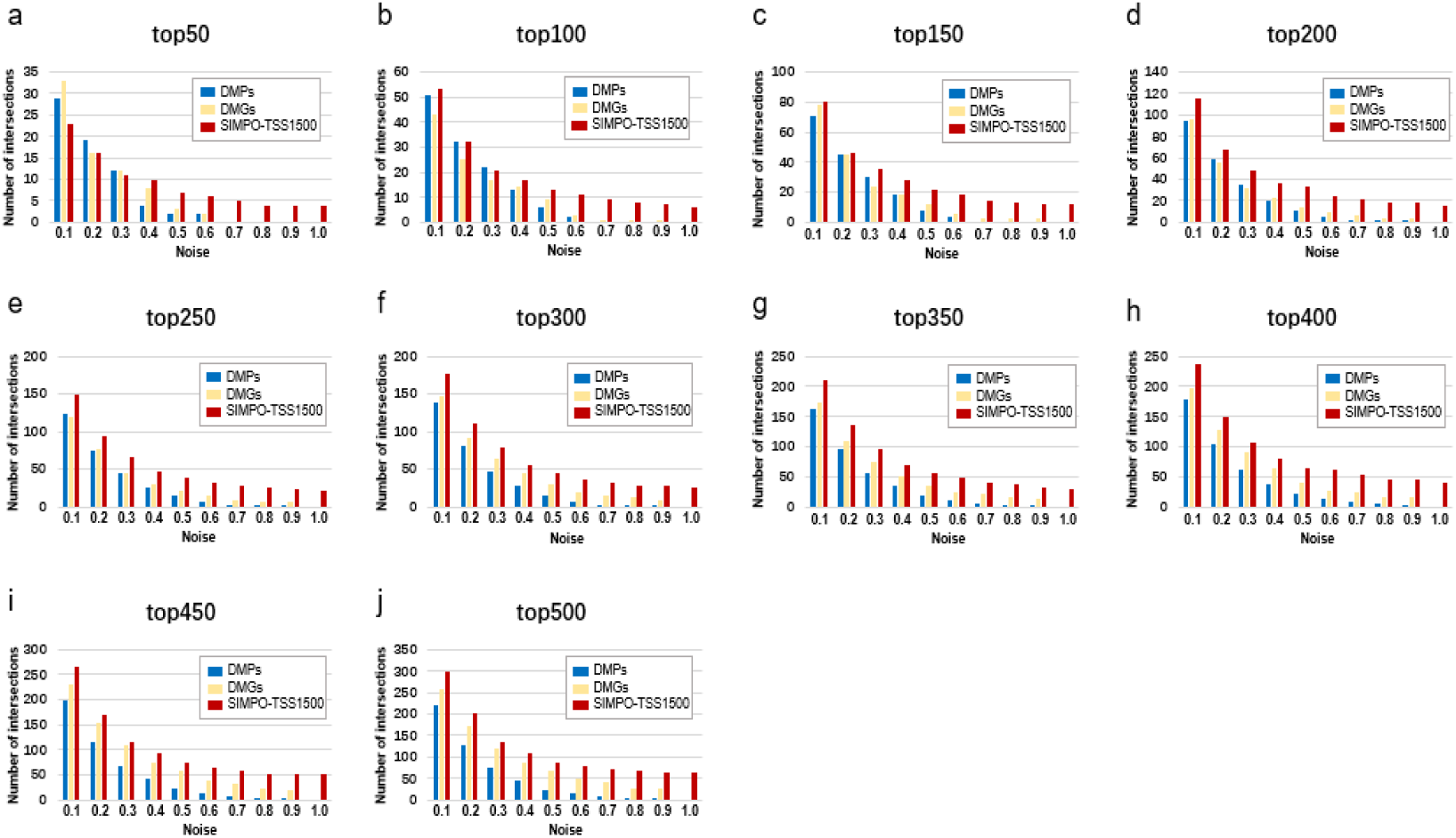
The intersections of top N smoking-associated genes identified by the original data and after adding noise-data of the GSE85210 dataset. The blue bars represent genes identified by the DMPs, the yellow bars represent genes identified by the DMGs, and the red bars represent genes identified by the SIMPO algorithm. (a) for top 50 probes/genes. (b) for top 100 probes/genes. (c) for top 150 probes/genes. (d) for top 200 probes/genes. (e) for top 250 probes/genes. (f) for top 300 probes/genes. (j) for top 350 probes/genes. (h) for top 400 probes/genes. (i) for top 450 probes/genes. (j) for top 500 probes/genes.

### 3.3 Robustness verification of SIMPO algorithm for different datasets

For traditional probe-based methods, because the same gene often contains multiple methylation probes, the same gene will get multiple smoking-related P-values. For the same gene, this study intended to select the average, the max, and the min P-value to represent the correlation between this gene and the smoking phenotype. The intersections of top N genes between dataset pairs were then used to show the robustness of the traditional method between different datasets. The results are shown in **Figure 8**: The min P-value probe-selected method is the most robust among different datasets. However, the proportion of intersections is relatively small at only about 5%.

**Figure 8.**
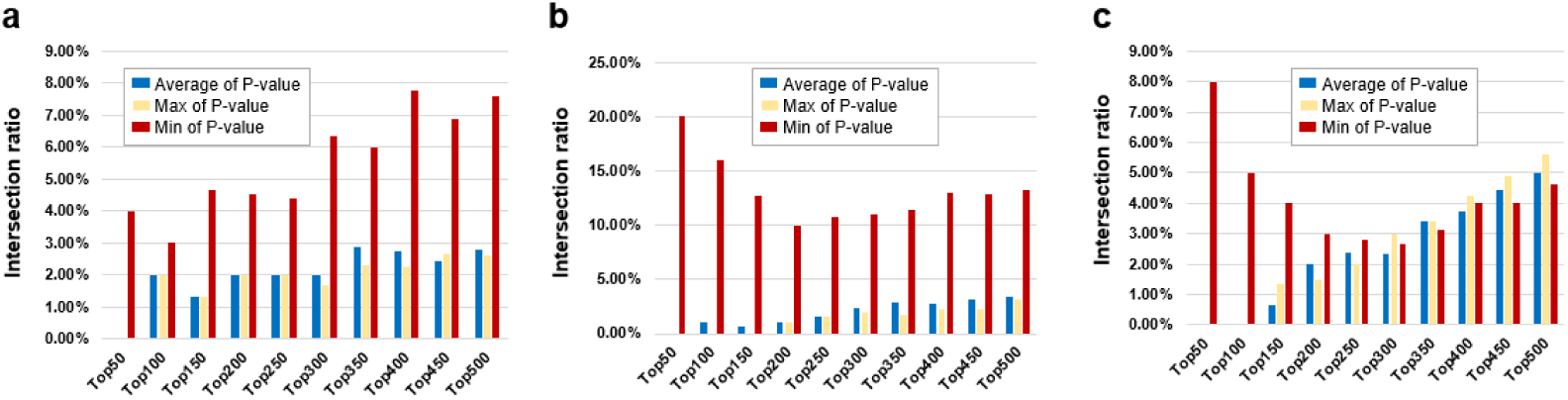
The intersection ratios of top N differential genes derived from different probes between different datasets (between the average, max, and min P-value of probes). The blue bars represent top N genes derived from the average P-value of probes, the yellow bars represent top N genes derived from max P-value of probes, and the red bars represent top N genes derived from min P-value of probes. (a) between GSE50660 and GSE53045 dataset. (b) between GSE50660 and GSE85210 dataset. (c) between GSE53045 and GSE85210 dataset.

Comparing the intersection ratio of differential genes/probes derived from at least two smoking-related datasets identified by DMGs and SIMPO showed that the robustness of the SIMPO algorithm (including SIMPO-TSS200, SIMPO-TSS1500, SIMPO-TSS200 & TSS1500) was significantly due to traditional EWAS (**Figure 9**). In the analysis of the Top N smoking phenotype-related genes, the SIMPO algorithm also obtained better results than the traditional probe-based method (DMPs) (**Figure 10**). In other words, the intersection ratios of smoking-associated genes identified by SIMPO in the two datasets were significantly higher than the DMPs.

**Figure 9.**
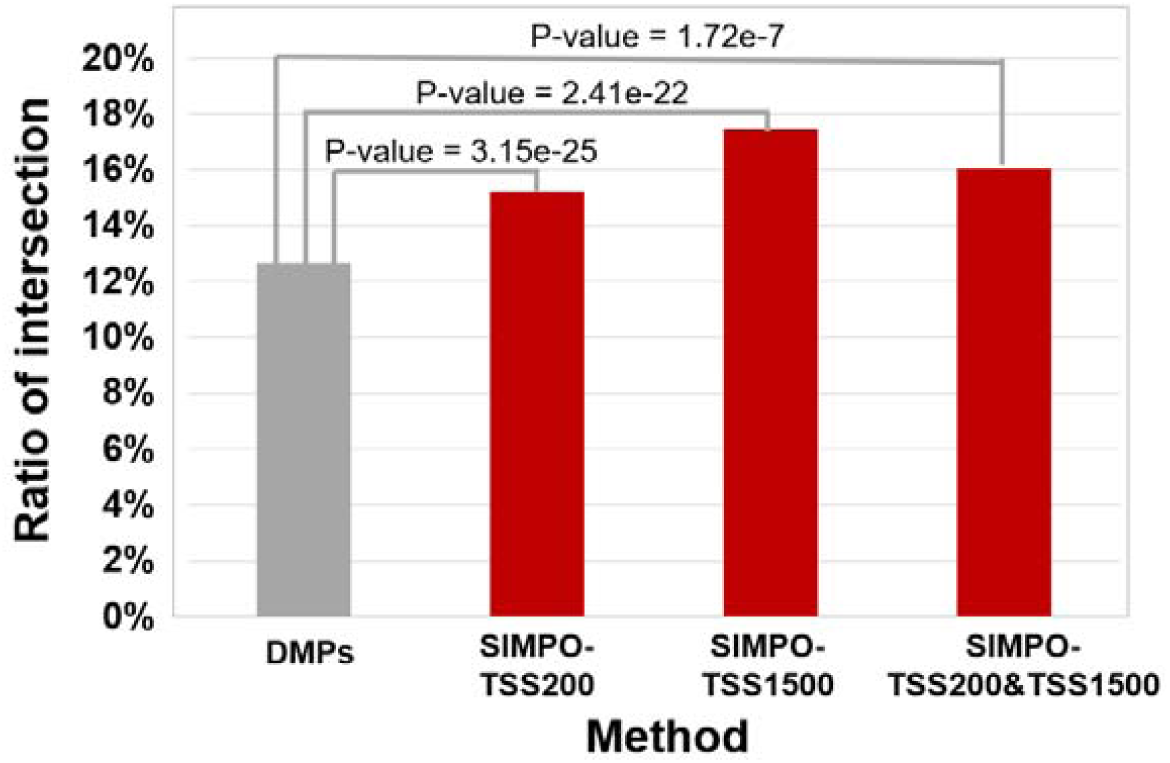
Comparison of the intersection ratio of differential genes/probes derived from at least two smoking-related datasets identified by different methods. The P-values were calculated by Chi-Square Test.

**Figure 10.**
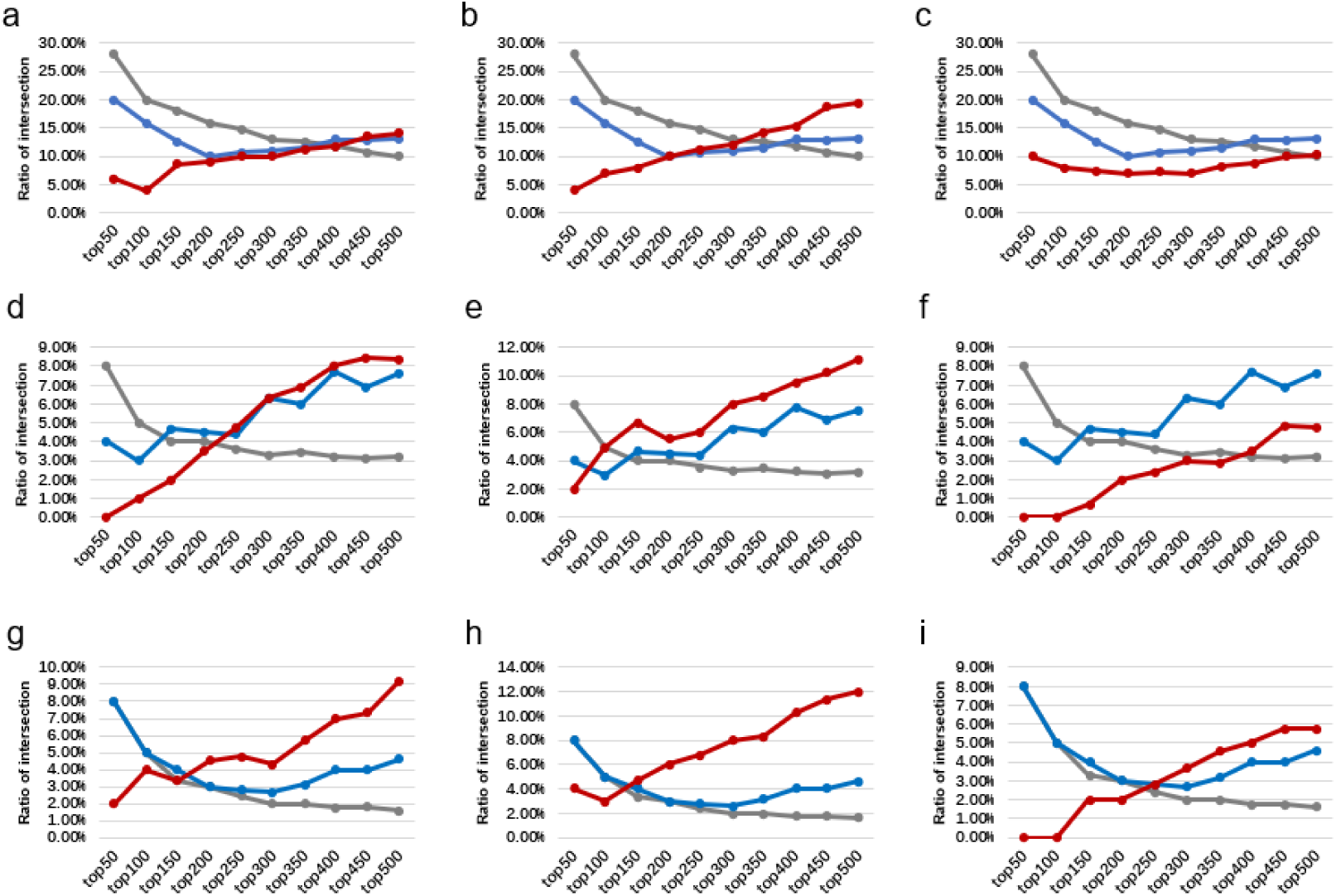
The intersection ratios of top N differential genes/probes between three smoking-related datasets. The gray, blue, and red lines represent the DMPs, DMGs, and SIMPO algorithm methods, respectively. (a) for SIMPO-TSS200 algorithm-identified differential genes between GSE50660 and GSE85210 dataset. (b) for SIMPO-TSS1500 algorithm-identified differential genes between GSE50660 and GSE85210 dataset. (c) for SIMPO-TSS200&1500 algorithm-identified differential genes between GSE50660 and GSE85210 dataset. (d) for SIMPO-TSS200 algorithm-identified differential genes between GSE50660 and GSE53045 dataset. (e) for SIMPO-TSS1500 algorithm-identified differential genes between GSE50660 and GSE53045 dataset. (f) for SIMPO-TSS200&1500 algorithm-identified differential genes between GSE50660 and GSE53045 dataset. (g) for SIMPO-TSS200 algorithm-identified differential genes between GSE53045 and GSE85210 dataset. (h) for SIMPO-TSS1500 algorithm-identified differential genes between GSE53045 and GSE85210 dataset. (i) for SIMPO-TSS200&1500 algorithm-identified differential genes between GSE53045 and GSE85210 dataset.

### 3.4 Biological significance verification of SIMPO algorithm

In this study, we verified the biological significance of the SIMPO algorithm by comparing the intersection of known tobacco use disorder-related genes (obtained from SCG-Drug database) with smoking phenotype-related genes that were identified by the traditional probe-based method (DMGs) and the SIMPO algorithm. The results are shown in **Table 2-4**: In the GSE50660 dataset, the proportion of known tobacco use disorder-related genes obtained by the SIMPO-TSS1500 and SIMPO-TSS200&TSS1500 algorithms is significantly better than DMGs; in the GSE53045 dataset, SIMPO-TSS1500 and the SIMPO-TSS200&TSS1500 algorithm is significantly better than DMGs; in the GSE85210 dataset, the SIMPO-TSS1500 algorithm is significantly better than DMGs (**Table S5-S8**).

**Table 1.**
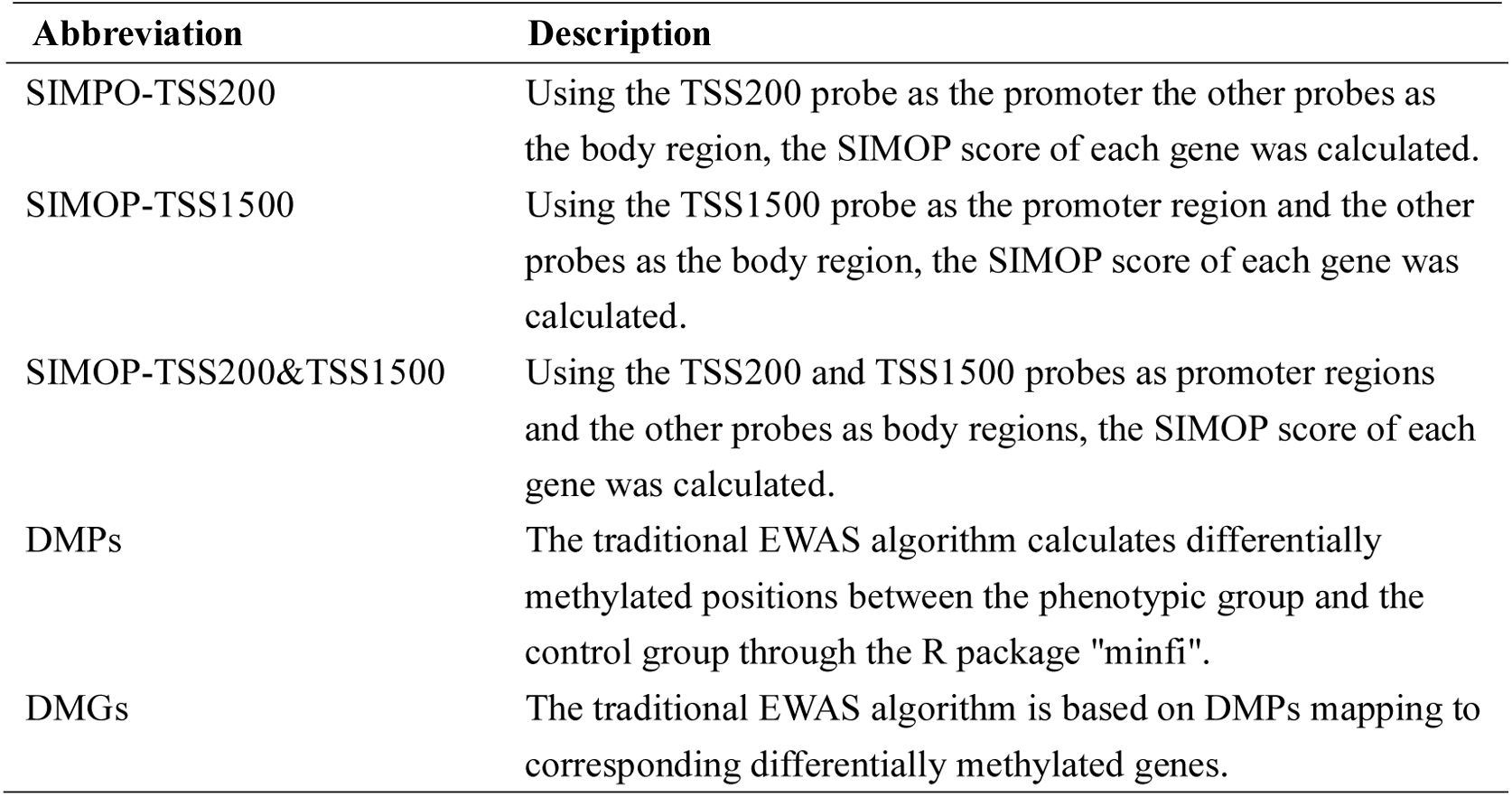
The description of the SIMPO algorithm and the traditional method.

| Abbreviation | Description |
| --- | --- |
| SIMPO-TSS200 | Using the TSS200 probe as the promoter the other probes as the body region, the SIMOP score of each gene was calculated. |
| SIMOP-TSS1500 | Using the TSS1500 probe as the promoter region and the other probes as the body region, the SIMOP score of each gene was calculated. |
| SIMOP-TSS200&TSS1500 | Using the TSS200 and TSS1500 probes as promoter regions and the other probes as body regions, the SIMOP score of each gene was calculated. |
| DMPs | The traditional EWAS algorithm calculates differentially methylated positions between the phenotypic group and the control group through the R package "minfi". |
| DMGs | The traditional EWAS algorithm is based on DMPs mapping to corresponding differentially methylated genes. |

**Table 2.** The known tobacco use disorder-related genes enriched by various algorithms of the GSE50660 dataset.

| Method | Background gene number | Intersection | Ratio | P-value<sup>1</sup> |
| --- | --- | --- | --- | --- |
| DMGs | 15147 | 2893 | 19.10% | \ |
| SIMPO-TSS200 | 827 | 168 | 20.31% | 3.87E-01<sup>2</sup> |
| SIMPO-TSS1500 | 705 | 156 | 22.13% | 4.61E-02<sup>2</sup> |
| SIMPO-TSS200&TSS1500 | 1639 | 383 | 23.37% | 3.45E-05<sup>2</sup> |

<sup>1</sup>calculated by Chi-Square Test.
<sup>2</sup>Compared to DMPs.

**Table 3.**
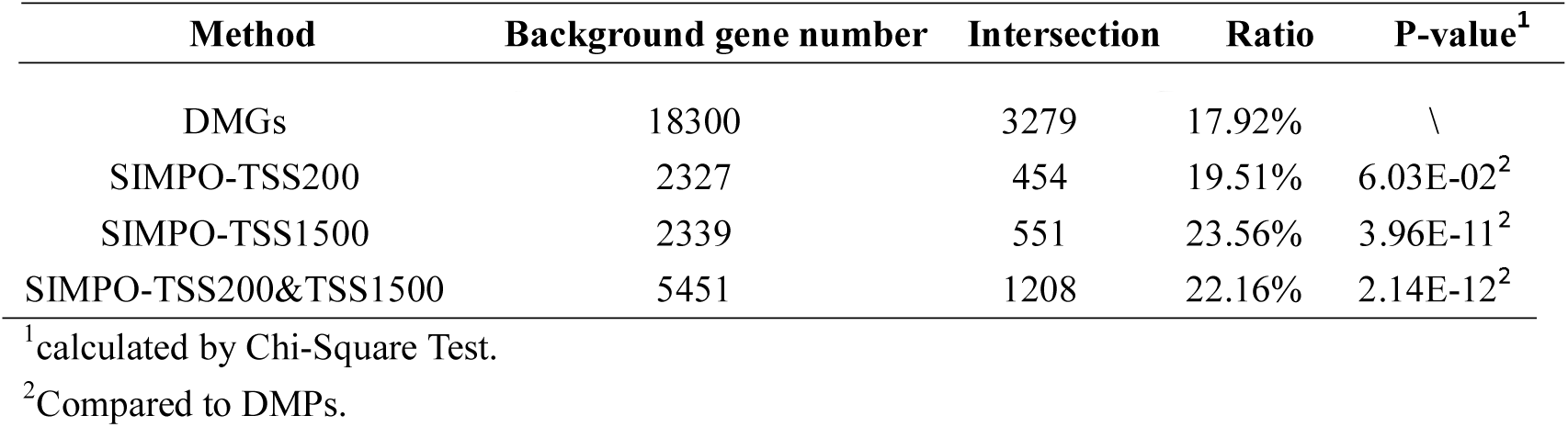
The known tobacco use disorder-related genes enriched by various algorithms of the GSE53045 dataset.

| Method | Background gene number | Intersection | Ratio | P-value<sup>1</sup> |
| --- | --- | --- | --- | --- |
| DMGs | 18300 | 3279 | 17.92% | \ |
| SIMPO-TSS200 | 2327 | 454 | 19.51% | 6.03E-02<sup>2</sup> |
| SIMPO-TSS1500 | 2339 | 551 | 23.56% | 3.96E-11<sup>2</sup> |
| SIMPO-TSS200&TSS1500 | 5451 | 1208 | 22.16% | 2.14E-12<sup>2</sup> |

<sup>1</sup>calculated by Chi-Square Test.
<sup>2</sup>Compared to DMPs.

**Table 4.** The known tobacco use disorder-related genes enriched by various algorithms of the GSE85210 dataset.

| Method | Background gene number | Intersection | Ratio | P-value<sup>1</sup> |
| --- | --- | --- | --- | --- |
| DMGs | 16846 | 3152 | 18.71% | \ |
| SIMPO-TSS200 | 790 | 155 | 19.62% | 5.22E-01<sup>2</sup> |
| SIMPO-TSS1500 | 1011 | 225 | 22.26% | 5.18E-03<sup>2</sup> |
| SIMPO-TSS200&TSS1500 | 1984 | 405 | 20.41% | 6.69E-02<sup>2</sup> |

<sup>1</sup>calculated by Chi-Square Test.
<sup>2</sup>Compared to DMPs.

In the above analyses, we analyzed a set of samples (including 1,202 individuals) that contained both transcriptome and DNA methylation data and showed that the SIMPO score of ∼40% of genes were significantly correlated with the average mRNA expression value, proving that SIMPO score and mRNA value of genes have good correlations. Next, we used three smoking-related DNA methylation datasets to validate the robustness of the SIMPO algorithm. The results showed that the robustness of the SIMPO is significantly better than the traditional probe-based methods for the same datasets and between different datasets. Finally, by comparing with known tobacco use disorder-associated genes, it is proved that the biological significance of the differential genes identified by the SIMPO algorithm is significantly superior to traditional methods, especially SIMPO-TSS1500. In summary, the SIMPO algorithm has good robustness and biological significance and can be further applied to related research in the field of epigenetic biology.

### 3.5 Application of SIMPO algorithm in IR-associated gene prediction

In this study, the SIMPO-TSS1500 algorithm was used to mining gene-level methylation remodeling pattern for IR-related dataset (GEO accession: GSE115278), and then *t-*test was applied to identify differential genes between individuals with HOMA-IR□≤□3 and >□3. As a result, 990 IR-associated genes were predicted by SIMPO-TSS1500 (**Table S9**). On the other hand, starting from the same dataset, another study has identified a total of 478 CpGs based on the traditional method, covering 499 differential genes [34]. Because IR is a pathological condition in which cells fail to respond appropriately to insulin, and it is a hallmark of type 2 diabetes [34,38], we speculated that above IR-related differential genes are associated with diabetes. By querying the known diabetes-associated genes recorded in the SCG-Drug database, it was found that only 77 genes of the 499 genes (15.43%) identified by traditional methods were known as diabetes-associated genes. For the 990 genes identified by the SIMPO-TSS1500 algorithm, the ratio is 44.44% (440 of 990 genes) (**Table S9**), which is significantly higher than the traditional method (P-value = 1.20e-28, based on chi-square test).

Then, according to the P-values of the differential genes obtained by the *t*-test, from small P-value (most significant) to large P-value (least significant), we obtained the top 100 ∼ top 1000 gene sets related to IR. Similarly, through the probe-based method, we also obtained the top 100 ∼ top 1000 gene sets. It is worth reminding that when a gene corresponds to multiple probes, we used the probe with the smallest P-value to represent this gene and to rank. Based on the KEGG pathway enrichment of the Enrichr database (http://amp.pharm.mssm.edu/Enrichr/), the results showed that multiple top N gene sets identified by SIMPO-TSS1500 were enriched in diabetes-related KEGG pathways (**Table 5**), while the top N gene sets identified by probe-based methods were not enriched in corresponding pathways.

**Table 5.** The diabetes-related KEGG pathway enrichment of SIMPO-predicted the top N gene sets.

| Diabetes-related KEGG pathway | Enriched gene set |
| --- | --- |
| Cell cycle | SIMPO-TSS1500 Top900; SIMPO-TSS1500 Top1000 |
| Maturity onset diabetes of the young | SIMPO-TSS1500 Top700; SIMPO-TSS1500 Top800;<br>SIMPO-TSS1500 Top900; SIMPO-TSS1500 Top1000 |
| Neurotrophin signaling pathway | SIMPO-TSS1500 Top600 |
| P53 signaling pathway | SIMPO-TSS1500 Top300; SIMPO-TSS1500 Top400;<br>SIMPO-TSS1500 Top500; SIMPO-TSS1500 Top600;<br>SIMPO-TSS1500 Top700; SIMPO-TSS1500 Top800;<br>SIMPO-TSS1500 Top900; SIMPO-TSS1500 Top1000 |
| Wnt signaling pathway | SIMPO-TSS1500 Top100; SIMPO-TSS1500 Top200;<br>SIMPO-TSS1500 Top300; SIMPO-TSS1500 Top400;<br>SIMPO-TSS1500 Top500; SIMPO-TSS1500 Top800;<br>SIMPO-TSS1500 Top900; SIMPO-TSS1500 Top1000 |

<sup>1</sup>obtained from Enrichr database (http://amp.pharm.mssm.edu/Enrichr/).

In addition, we also conducted disease enrichment for the IR-associated gene sets predicted by SIMPO and DMGs-based methods. The results are shown in **Figure 11**, SIMPO-predicted top N gene sets were enriched to a variety of diabetes-related diseases through Enrichr database, including non-insulin-dependent diabetes mellitus, permanent neonatal diabetes mellitus, maturity onset diabetes mellitus in young, neonatal diabetes mellitus, and obtained 15 gene sets-disease associations. DMGs-predicted top N gene sets only obtained 9 such associations. In summary, the results show that the biological significance of IR-associated genes predicted by SIMPO-TSS1500 is better than those predicted by DMGs-based methods.

**Figure 11.**
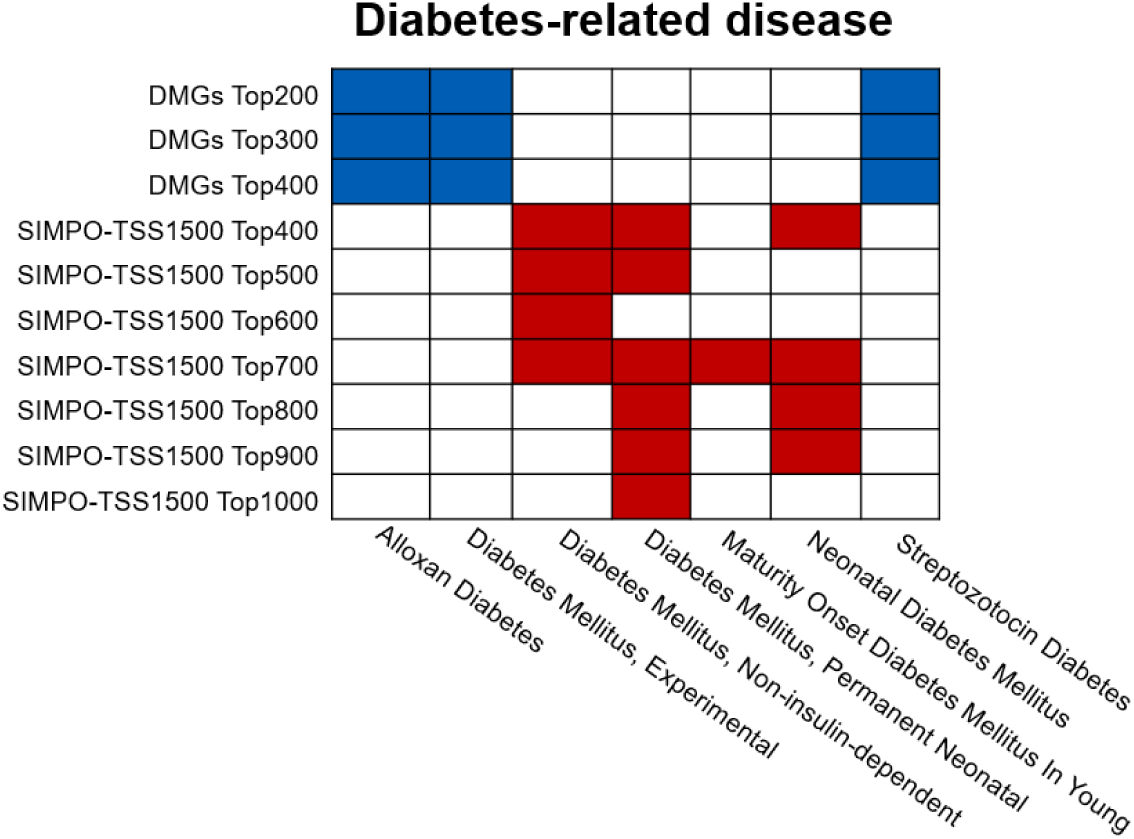
The diabetes-related disease enrichment of DMGs- and SIMPO-predicted top N gene sets. Enrichment analysis was obtained from the Enrichr database (http://amp.pharm.mssm.edu/Enrichr/). The diabetes-related diseases, including alloxan diabetes, experimental diabetes mellitus, non-insulin-dependent diabetes mellitus, permanent neonatal diabetes mellitus, maturity onset diabetes mellitus in young, neonatal diabetes mellitus, and streptozotocin diabetes. The blue squares indicate that the DMGs-predicted top N gene set is significantly associated with the corresponding diabetes-associated disease; the red squares indicate that the SIMPO-predicted top N gene set is significantly associated with the corresponding diabetes-related disease.

### 3.6 Application of SIMPO algorithm in PD-associated gene prediction

SIMPO-TSS1500 algorithm was further used to mining gene-level methylation remodeling of PD patients and control individuals. Then, there were 959 significantly differential genes for the GSE72774 dataset (**Table S10**), and 1,077 significantly differential genes for the GSE111629 dataset related to PD have been identified by a *t*-test (**Table S10**). In addition, combining the above two DNA methylation datasets, previous researchers predicted a total of 82 PD-related significant difference CpGs based on the traditional EWAS method, corresponding to 62 genes [27]. By querying the known PD-associated genes in SCG-Drug database, it was found that only 4 of 62 genes (6.45%) identified by the traditional method were known PD-associated genes. For the SIMPO-TSS1500-identified PD-associated genes, the ratios were 9.19% (for GSE111629) and 9.28% (for GSE72774) (**Table S10**), which are higher than the traditional methods.

Then, this study enriched the KEGG pathway for SIMPO-TSS1500-predicted differential gene sets of PDs through GSEA. The results are shown in **Table 6**: These two PD-related gene sets were enriched to 12 KEGG pathways. By querying the biological function annotations for the pathways on the KEGG website (https://www.genome.jp/kegg/pathway.html), it was found that four pathways are related to nervous system diseases, including Alzheimer’s disease, Inositol phosphate metabolism, phosphatidylinositol signaling system, and purine metabolism.

**Table 6.** The KEGG pathway enrichment of SIMPO-predicted PD-associated genes.

| Dataset | KEGG pathway | Annotation |
| --- | --- | --- |
| GSE72774 | Alzheimers disease | Nervous system diseases-related |
| GSE72774 | Cysteine and methionine metabolism | \ |
| GSE72774, GSE111629 | Cytokine cytokine receptor interaction | \ |
| GSE111629 | Glycolysis gluconeogenesis | \ |
| GSE72774, GSE111629 | Gnrh signaling pathway | \ |
| GSE72774, GSE111629 | Inositol phosphate metabolism | Nervous system diseases-related |
| GSE72774 | Jak stat signaling pathway | \ |
| GSE111629 | Lysosome | \ |
| GSE111629 | Phosphatidylinositol signaling system | Nervous system diseases-related |
| GSE72774 | Purine metabolism | Nervous system diseases-related |
| GSE72774 | Snare interactions in vesicular transport | \ |
| GSE111629 | Viral myocarditis | \ |

<sup>1</sup>calculated by GSEA (Gene Set Enrichment Analysis).

SCG-Drug and DisGeNET databases collected gene-disease associations from multiple sources, and both annotated the credibility scores of gene-disease associations. This study compared SIMPO-TSS1500 predicted PD-related differential gene sets with the top 10% scored PD pathogenic genes recorded in DisGeNET and SCG-Drug. The intersections of SIMPO-TSS1500 predicted gene sets with known PD-causing genes were significantly higher than the background databases (**Table 7**) (**Table S10**). The above results further proved the reliability of the SIMPO-predicted PD-associated gene sets. Moreover, it also reflects the robustness of mining the statistical difference of DNA methylation between the promoter and other regions to identify gene-level associations with a given phenotype (SIMPO algorithm) from the side.

**Table 7.** Comparison of SIMPO-predicted with known PD-associated genes.

| Dataset | PD gene source | SIMPO-derived ratio | Background ratio | P-Value<sup>1</sup> |
| --- | --- | --- | --- | --- |
| GSE111629 | SCG-Drug<sup>2</sup> | 9.19% (99/1077) | 7.21% (351/4868) | 3.26E-03 |
|  | DisGeNET<sup>3</sup> | 7.15% (77/1077) | 5.88% (286/4868) | 2.79E-02 |
| GSE72774 | SCG-Drug<sup>2</sup> | 9.28% (89/959) | 7.21% (351/4868) | 4.27E-03 |
|  | DisGeNET<sup>3</sup> | 7.40% (71/959) | 5.88% (286/4868) | 1.68E-02 |

<sup>1</sup>calculated by Hypergeometric test.
<sup>2</sup>known PD-associated genes were collected from SCG-Drug (http://zhanglab.hzau.edu.cn/scgdrug).
<sup>3</sup>known PD-associated genes were collected from DisGeNET (http://www.disgenet.org).

## 4. Discussion

In recent years, through EWAS, researchers have identified thousands of phenotype-related differential methylation sites. However, since the same gene may contain hundreds of methylation sites, the DNA methylation beta values at different sites vary widely. Furthermore, DNA methylation remodeling has a certain degree of randomness on the genome. As a result, for multiple EWASs focused on the same phenotype, the identified differential methylation site intersections are small, which makes it challenging to identify phenotype-associated genes and analyze epigenetic mechanisms. Therefore, how to integrate the methylation values of different sites on the same gene and identify robust gene-level associations with phenotype becomes a challenge in the epigenetics field.

In this study, by analyzing a set of individual samples containing both transcriptome and DNA methylome data, it was found that the SIMPO scores of ∼ 40% genes were significantly correlated with average mRNA transcription, demonstrating that the SIMPO scores of genes have a reasonable correlation with gene expression. Then, three DNA methylation datasets related to the smoking phenotype were used to test the robustness of SIMPO algorithm. The results showed that the robustness of the SIMPO algorithm in the same dataset and between different datasets were significantly better than the traditional EWAS method. Finally, through comparing with known tobacco use disorder pathogenic genes, it is proved that the biological significance of SIMPO algorithm is also better than that of the traditional EWAS method, notably the SIMPO-TSS1500 algorithm. In summary, the SIMPO algorithm has good robustness and biological significance and can be further applied to related research in the field of epibiology. Therefore, we further applied the SIMPO algorithm to predict IR- and PD-associated genes and proved the biological significance of corresponding genes.

However, the SIMPO algorithm still has some shortcomings. In order to ensure the stability of the SIMPO algorithm, it requires that the promoter region and other regions of a gene contain five or more probes to obtain a SIMPO score. Therefore, a large number of genes containing a small number of probes will be lost. At present, the number of genes that can be calculated by SIMPO-TSS200 and TSS1500-SIMPO is only about 4,000, and the number of genes that can be calculated by SIMPO-TSS200 & 1500 is only about 10,000. It is much smaller than the number of genes contained in the human genome. As a result, some critical functional genes have been missed for the current SIMPO algorithm. Fortunately, with the popularity of the Illumina 850K BeadChip in EWAS, which contains more than 850,000 probes, the number of genes that the SIMPO algorithm can calculate will increase significantly.

## 5. Ethics Statement

No.

## 7. Author Contributions

Yuan Quan and Fengji Liang conducted the data mining and bioinformatics analyses; Jianghui Xiong led the epigenetic research, and designed the strategy for integrated analysis of DNA methylation data; Yuexing Zhu and Ying Chen took part in the data analysis; Ruifeng Xu helped in preparing the manuscript; Yuan Quan and Jianghui Xiong wrote the manuscript.

## 9. Availability of data and materials

The datasets supporting the results of this article are included within the article or in additional files.

